# Effects of Fecal Source Input, Environmental Conditions, and Environmental Sources on Enterococci Concentrations in a Coastal Ecosystem

**DOI:** 10.1101/311928

**Authors:** Derek Rothenheber, Stephen Jones

## Abstract

Fecal pollution at coastal beaches in the Northeast, USA requires management efforts to address public health and economic concerns. Concentrations of fecal-borne bacteria are influenced by different fecal sources, environmental conditions, and ecosystem reservoirs, making their public health significance convoluted. In this study, we sought to delineate the influences of these factors on enterococci concentrations in southern Maine coastal recreational waters. Weekly water samples and water quality measurements were conducted at freshwater, estuarine, and marine beach sites from June through September 2016. Samples were analyzed for total and particle-associated enterococci concentrations, total suspended solids, and microbial source tracking markers for multiple sources. Water, soil, sediment, and marine sediment samples were also subjected to 16S rRNA sequencing and SourceTracker analysis to determine the influence from these environmental reservoirs on water sample microbial communities. Enterococci and particle-associated enterococci concentrations were elevated in freshwater, but suspended solids concentrations were relatively similar. Mammal fecal contamination was significantly elevated in the estuary, with human and bird fecal contaminant levels similar between sites. A partial least squares regression model indicated particle-associated enterococci and mammal marker concentrations had the most significant positive relationships with enterococci concentrations across marine, estuary, and freshwater environments. Freshwater microbial communities were significantly influenced by underlying sediment while estuarine/marine beach communities were influenced by freshwater, high tide height, and estuarine sediment. We found elevated enterococci levels are reflective of a combination of increased fecal source input, environmental sources, and environmental conditions, highlighting the need for encompassing MST approaches for managing water quality issues.

**IMPORTANCE:** Enterococci have long been the federal standard in determining water quality at estuarine and marine environments. Although enterococci are highly abundant in the fecal tracts of many animals they are not exclusive to that environment and can persist and grow outside of fecal tracts. This presents a management problem for areas that are largely impaired by non-point source contamination, as fecal sources might not be the root cause of contamination. This study employed different microbial source tracking methods to delineate influences from fecal source input, environmental sources, and environmental conditions to determine which combination of variables are influencing enterococci concentrations in recreational waters at a historically impaired coastal town. Results showed that fecal source input, environmental sources and conditions all play a role in influencing enterococci concentrations. This highlights the need to include an encompassing microbial source tracking approach to assess the effects of all important variables on enterococci concentrations.

## INTRODUCTION

Fecal contamination of coastal recreational waters is a significant public health concern, as fecal material, often from nonpoint sources, can harbor an array of different pathogens. The US EPA has established regulations based on enterococci bacteria as the indicator of fecal-borne pollution to help manage water quality at estuarine and marine beaches (1). These organisms correlated well with predicted public health outcomes in several epidemiological studies that served as the basis for their adoption as the regulatory water quality indicator (2–5). The presence of human feces can present an elevated public health risk in recreational waters compared to non-human sources due to the lack of an “inter-species barrier” for diseases and the higher density of human pathogens that humans can carry (6–8). Although human pollution represents the greatest public health risk, other fecal sources that contain enterococci and possibly human pathogens can be chronic or intermittent sources of both, making beach water quality management and remediation efforts more complex.

The need to differentiate fecal sources in recreational waters led to the emergence of microbial source tracking (MST) methods in the early 2000s, most notably the PCR-based assays that target the 16S rRNA gene in *Bacteroides* spp. (9, 10). There are a wide range of species-specific genetic markers designed to identify human fecal sources and various domestic and wildlife fecal sources. These assays have been in use for well over a decade and are supported by numerous and rigorous laboratory evaluations and field applications (11–17). Initial field studies investigated the relationship between MST markers and FIB concentrations in recreational waters to better elucidate potential sources of fecal pollution. Some studies have found strong relationships between the MST markers and enterococci (12, 18) while other studies have found either weak or no relationships (19–21), many of which are discussed in a review by Harwood et al. (22). One main factor affecting the relationship between enterococci and the relative strength of different sources of fecal contamination is that enterococci can persist and grow in the environment, which can significantly influence their concentrations in recreational water (23).

Due to the pervasiveness of enterococci in natural ecosystems, recent studies have been conducted to not only elucidate environmental parameters controlling their growth, but also to identify naturalized niches that can act as reservoirs for enterococci and the associated influence on water quality measurements. Specifically, enterococci have been shown to persist in fresh water sediments (24–26) and marine sediments (24, 27), and in some cases their relative concentrations in sediments are several orders of magnitude higher than the overlying water (24, 28–30). In addition, enterococci persist in soils affected by anthropogenic activities (31) as well as more natural soil environments (32–34). Thus, soil can act as a significant reservoir of enterococci that can, if eroded, confound concentrations observed in recreational waters. Evaluating the influence of sediment and or soil on water quality has, in some studies, been conducted by measuring total suspended solids as a surrogate for sediment-associated enterococci (27, 35, 36), however this non-specific approach does not indicate the specific type of source(s) of the suspended solids. With the advent of next generation sequencing, sources of sediment or soil bacteria can be fingerprinted via 16S rRNA sequencing, and programs like SourceTracker can then determine relative fractions of source-specific 16S fingerprints within a water sample (37).

This study examined the coastal and estuarine beaches of Wells, ME where there has been historically elevated enterococci levels, as reported by the Maine Healthy Beaches Program (38). Prior to this study, only a ribotyping-based MST study (39) that also involved other indicator tracking work had been conducted in this area. In that study, the two major freshwater inputs, the Webhannet River and Depot Brook were found to be the major influences on water quality related to an array of fecal contamination sources. To investigate potential sources of enterococci we measured three major categories of variables (fecal source input, environmental conditions, and environmental sources) and then used a partial least squares regression model approach to determine the most significant influences on the enterococci concentrations in water samples.

## RESULTS

### Total and particle-associated enterococci concentrations and total suspended solids in water

During this study, total enterococci concentrations were highest in freshwater sites, with concentrations significantly decreasing from there to the estuary and then the marine beach areas (Figure 2). The geometric mean enterococci concentrations were 197 and 40 CFU/100 ml at the Depot and Webhannet sites, respectively, with 71% of samples exceeding 104 CFU/100 ml at the Depot site compared to 21% at the Webhannet site. In contrast, the geometric mean enterococci concentrations at the other sites were all <15 CFU/100 ml and samples exceeded 104 CFU/100 ml 0% (at Wells Beach) to 25% of the time. In addition to measuring enterococci concentrations in water samples, particle-associated enterococci and suspended solid concentrations were measured to better understand the potential mode of transport of these bacteria within this coastal watershed. Throughout the study period (June-September 2016), levels of total and particle-associated enterococci varied by site. Concentrations were lowest at the marine beach (Wells Beach) compared to other sites, with levels significantly higher in all estuary sites (W11-W15) and freshwater sites (Depot & Webhannet; Figure 2).

**Figure 1:**
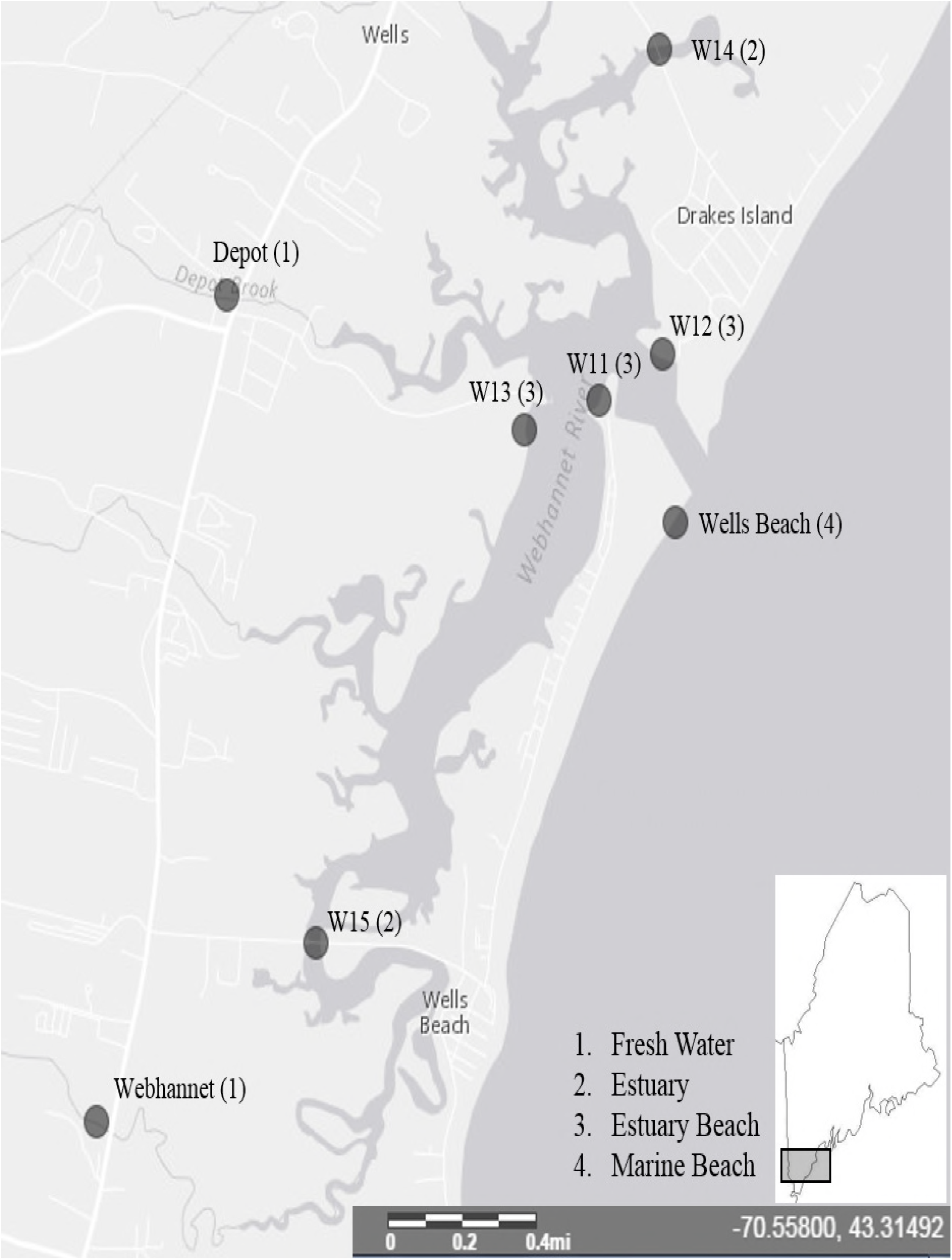
Wells Maine Study area and sampling sites. All water collection sites are marked with a dark grey circle. Sites that correspond to fresh water are indicated with a (1), estuary (2), estuary beach (3), and marine beach (4).

**Figure 2:**
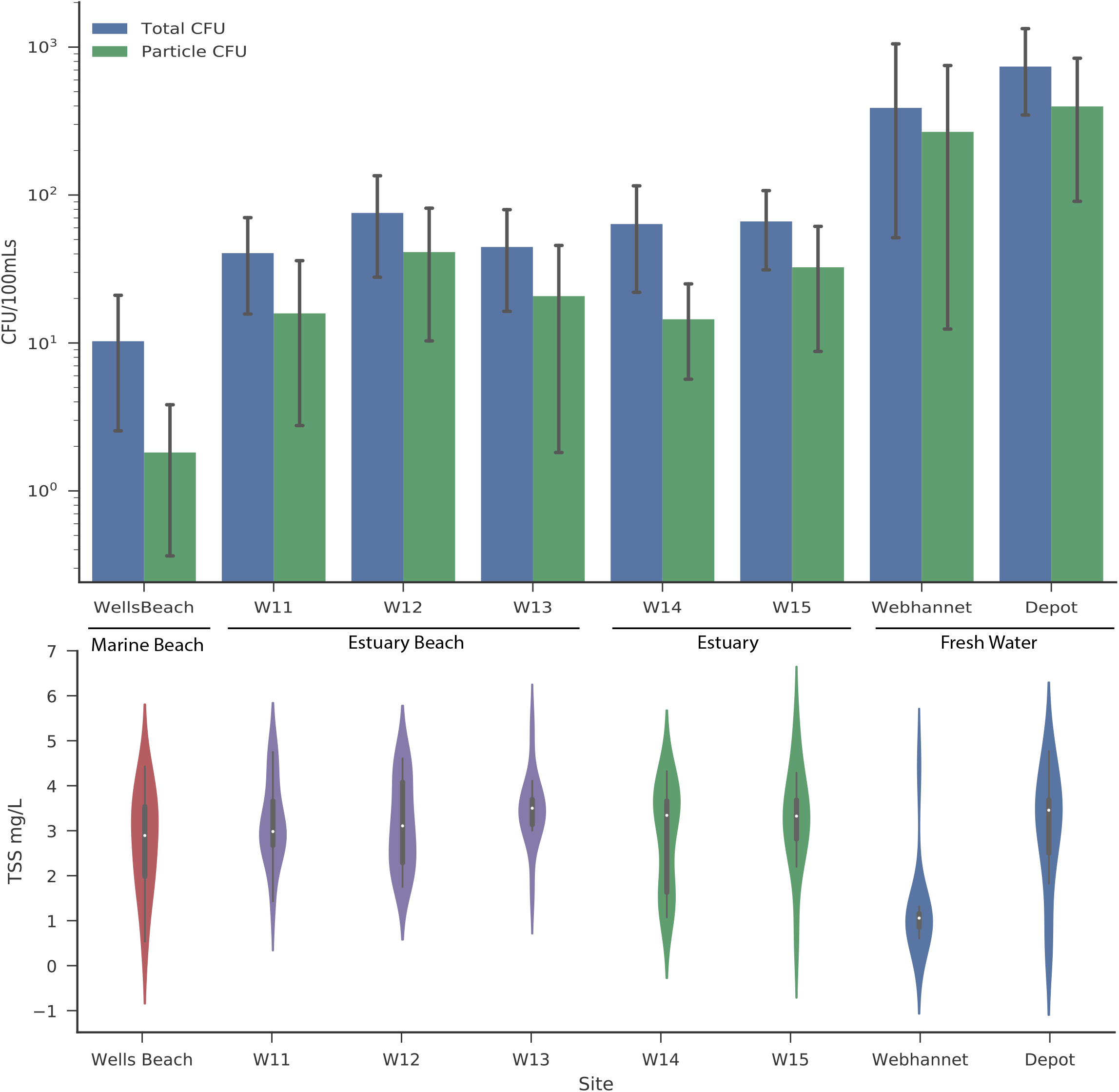
Geometric Mean Concentrations of Total and Particle Associated Enterococci and Average Total Suspended Solids Concentrations at the Eight Study Sites. (A) Total enterococci concentrations are represented with the blue bar, and particle associated enterococci concentrations correspond to the green bar. Error bars are derived from variation from each site across the entire study. (B) Violin plots were used to represent TSS concentrations, and the color corresponds to the type of site including marine beach (red), estuary beach (purple), estuary (green), or fresh water (blue). Horizontal lines go through the median of each violin plot.

Both total and particle-associated enterococci geometric mean concentrations were statistically similar at the estuary beach (W11, W12, W13) and estuary (W14, W15) sites. Freshwater sites (Webhannet and Depot) however, had statistically higher enterococci concentrations than other sites (Figure 2; p < 0.05). The ratio of total to particle-associated enterococci varied throughout the season, with an average of 36.3% (SD ± 30) across all sites. Sites within the estuary beach showed the highest ratio (41%, SD ± 32), however there were no significant differences observed between sites or types of sites. Average TSS concentrations were relatively low and similar for most sites, with an overall average of 2.9 mg TSS/L (SD± 1.2). The Webhannet freshwater site, however, had a significantly lower average TSS concentration (1.2 mg/L ± 1.0SD, p < 0.05) (Figure 2), despite, as previously mentioned, having higher enterococci concentrations. The relationship between particle-associated enterococci and TSS was not significant (r^2^ = 0.0011), and significant rainfall events were seldom and sparse with only one greater than 1 in 48 h prior to sampling. Overall, this study showed enterococci concentrations were significantly different by site and were ubiquitously associated with particles, which was independent of suspended solids concentrations.

### Presence of fecal sources in fresh, estuarine, and marine waters

The concentration of fecal pollution in this study area was determined using both PCR and quantitative PCR MST assays to identify and quantify predominant sources of fecal contamination present in the water. The mammal fecal marker (Bac32) was detected via PCR at all sites 100% of the time throughout the study period. (Supplementary Material 1E). The human fecal marker (HF183) was detected in 51% of all water samples, with the highest detection rate in fresh water (56%) and the lowest detection rate in marine beach water (46%). Differences in the percent detection of the gull fecal marker (Gull2) were most pronounced between freshwater (10%) and all other sites (>77%). The dog fecal marker (DF475) detection rate was highest in the estuary beach water (10/44 = 23%), however 8 of the 10 positive samples were detected in July (8/13 = 61%). For all other sites, an increase in the detection of dog fecal marker also occurred during July, with 44% (16/36) detection, compared to 0% for August and September and <1% for June. Thus, most of the dog contamination at all sites was associated with unknown dog-related conditions during July.

### Concentrations of mammal, human, and bird fecal sources

We used qPCR to provide relative quantitative measures of mammal, human and bird fecal contamination levels. Water at estuary and estuary beach sites contained significantly higher levels of mammal (AllBac) fecal marker copies, with an average of 1.54 × 10^7^ compared to 2.62 × 10^6^ in freshwater and 3.9 × 10^6^ copies/100 ml in marine beach (p < 0.05). Average concentrations of human (HF183) and bird (GFD) fecal markers were not statistically different between sites, however, concentrations of the human marker in individual samples varied from 0 – 2.04 × 10^4^ copies/100 ml (Figure 3), while bird fecal marker concentrations were relatively stable across all sites. No significant temporal trends were observed for any of the quantitative fecal marker levels. Compared with presence/absence detection of fecal sources, quantitative measurements also did not show strong spatial patterns, except mammal marker levels showed significant increases at estuary and estuary beach sites compared to marine and freshwater sites.

**Figure 3:**
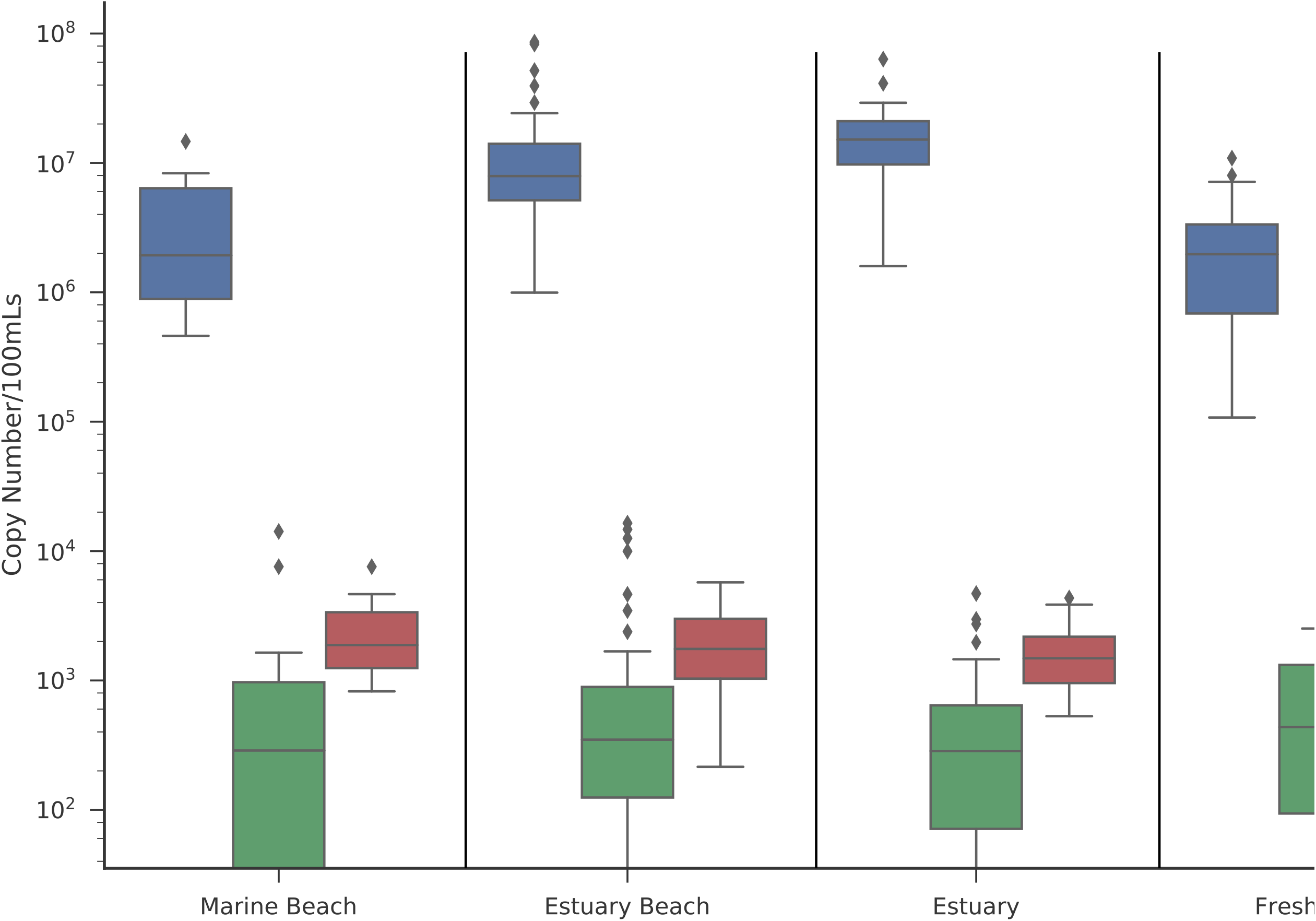
Relative Levels of Mammal, Human, and Bird Fecal Source at the Different Types of Study Sites. Box plots represent levels of microbial source tracking markers at marine beach (Wells Beach), estuary beach (W11, W12, W13), estuary (W14 & W15), and fresh water (Webhannet & Depot). Outlier data are represented with a black diamond.

### Differences between water, soil, and sediment bacterial community compositions

16S amplicon sequencing was used to characterize the microbial community present in water and other sample matrices (soil, sediment, and marine sediment), which was the nexus for ensuing SourceTracker analysis. A total of 3,276,196 reads and 7,706 unique OTUs were obtained from the 177 samples of fresh, estuary, estuary beach and marine beach water and soil, sediment, and marine sediment. The number of OTUs assigned and the Shannon diversity index were significantly higher for soil, sediment, and marine sediment when compared to water samples (Figure 4, p < 0.05). Most taxa in the estuary and marine beach water samples were identified as Flavobacteriia, Alphaproteobacteria, and Gammaproteobacteria classes, which together accounted for 84% of the total assigned taxa. Cyanobacteria accounted for 34% of the taxa in marine sediment, and Betaproteobacteria was one of the top three most abundant taxa in fresh water, soil and sediment (Figure 4). A Non-Metric Multi-Dimensional Scaling (NMDS) ordination was used to determine if the bacterial communities from water and other matrices (soil and sediments) differed based on their taxonomic composition. Bacterial communities from the marine beach and estuary (All Estuary) waters were similar, but were statistically different from fresh water (Figure 5, p < 0.05). The bacterial communities associated with soil, sediment and marine sediment were all distinct when compared to each other and water samples, indicating unique groups of OTUs (Figure 5, p < 0.05). Samples taken from different areas within the watershed (soil, estuarine water, freshwater, etc.) contained unique bacterial compositions, allowing for downstream analysis with the SourceTracker software to discern relative contributions of these different communities to the make-up of microbial communities in the different types of water samples.

**Figure 4:**
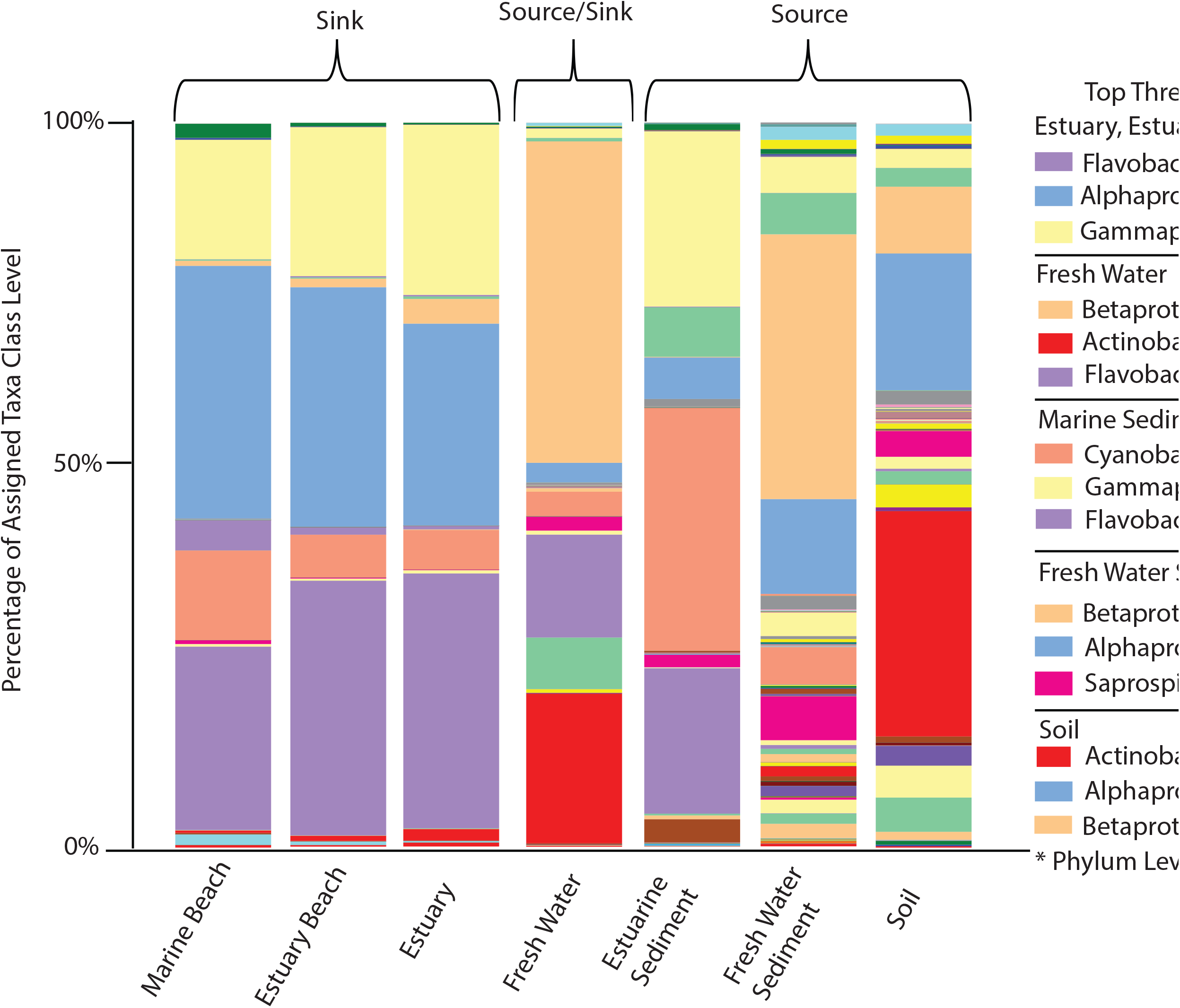
16S Taxa Profiles and the Top Three Most Abundant Bacterial Classes in All Source and Sink Samples. Stacked bar plots represent percentages of the class level composition of the microbial communities. Source corresponds to environmental sources that were finger-printed with the SourceTracker program, and then used to determine their presence within water (sink) samples. The table represents the top three classes for each group of samples and * corresponds to phylum level. For a complete list of all taxa assignments refer to Supplementary material 4.

**Figure 5.**
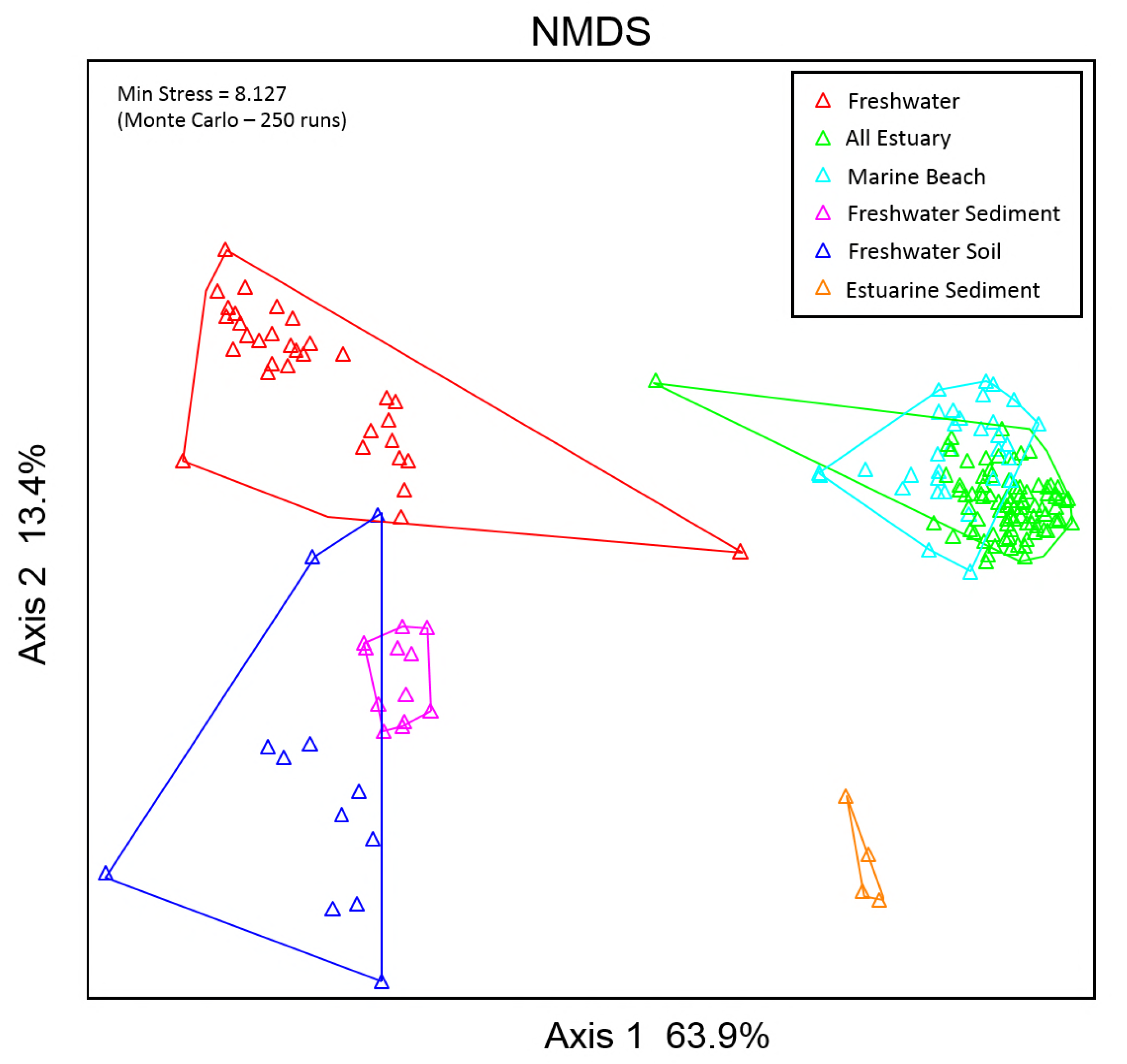
Differences Between Microbial Communities from Different Source Materials. Samples are color-coded based on sample matrix (i.e. soil, fresh water, etc.). Percent of variation explained are displayed on the x and y axis and the minimum stress of the ordination is shown in the top left corner.

### Environmental source contribution to water samples

The fraction of freshwater, sediment, soil, estuarine sediment, and marine beach water source bacterial communities within estuary and estuary beaches water samples were calculated using the Bayesian mixing model SourceTracker. Freshwater sample analysis showed a high probability of taxa originating from underlying sediment (74%) and much lower probability of taxa originating from soil (2.6%). Initial results for the estuary and estuary beach indicated that marine beach water was the dominant source of bacteria (Table 1). However, given that likely fecal sources are coming from the watershed, we excluded marine beach water as a potential source and included it as a sink then re-analyzed the data. These second results showed that freshwater taxa had a high probability of being a significant fraction of estuary (73%), estuary beach (66%) and marine beach (35%) water communities, with a significantly higher percentage for the estuary locations compared to the marine beach (Table 1, p < 0.05), which is more influenced by ocean microbial taxa. Despite the significant percentage of freshwater taxa assignments in the estuary, estuary beach, and marine beach waters there were no freshwater sediment or soil taxa assignments for these sites. The data for the percent of unidentifiable taxa showed the opposite trend compared to percent of assigned freshwater taxa. Unidentifiable taxa in the marine beach were significantly higher (46%; p < 0.05), which is not surprising given that marine beach water community would likely be most influenced by non-terrestrial sources. Estuarine sediment was the highest likely identified source in the water from the marine beach site (19%), and it was significantly higher than percentages calculated for all estuary sites (p < 0.05). Overall results showed that freshwater source-related taxa were a pervasive source throughout the estuary and marine beach, and while sediment source-related taxa were highly abundant in the freshwater they were not observed within the estuary or marine beach.

**Table 1.**
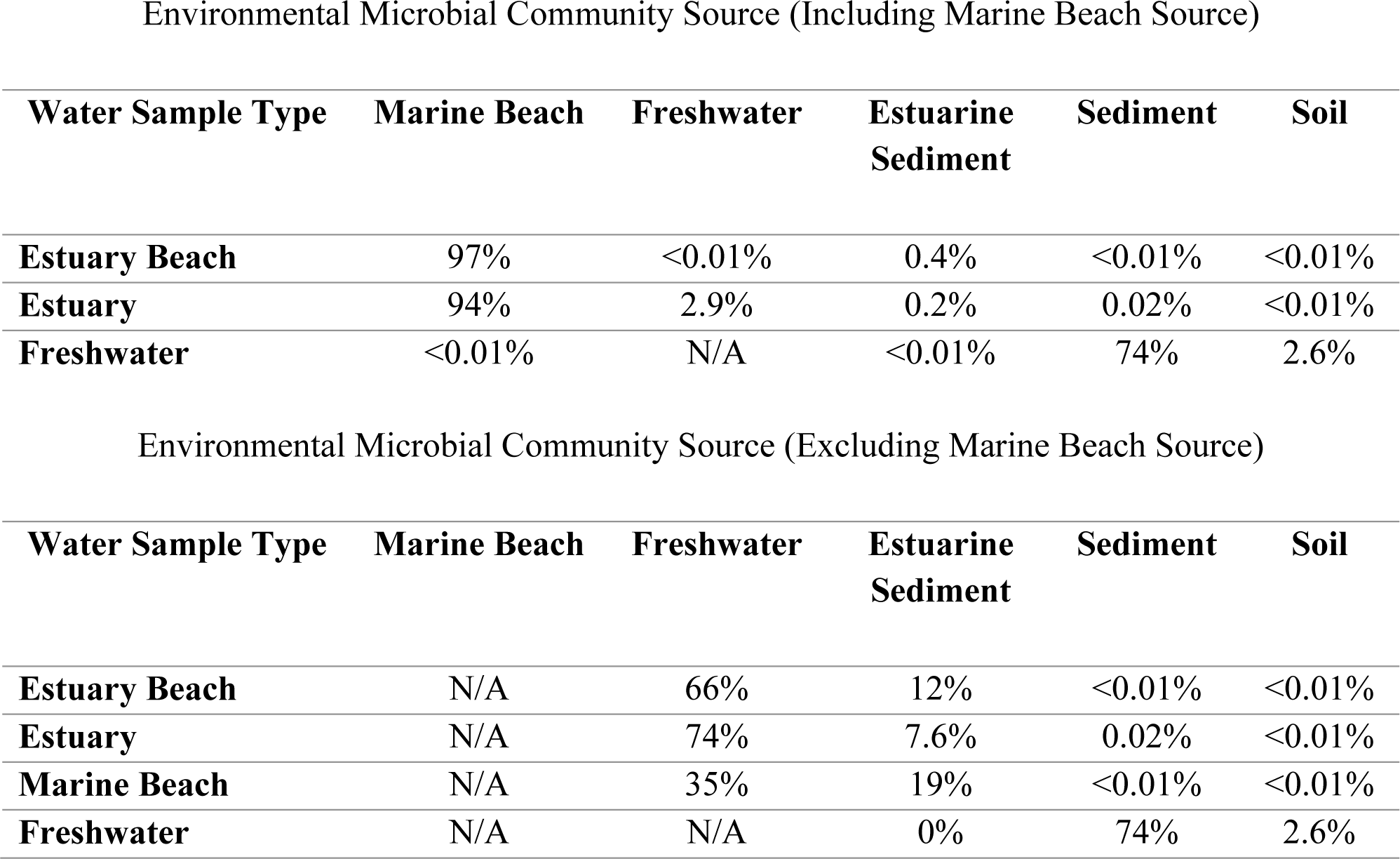
The relative contribution of different sources to the microbial communities in estuarine and marine water. SourceTracker was run with two different configurations, one where Marine Beach water was included as a potential source (top) and a second run where Marine Beach water was excluded as a potential source (bottom).

Environmental Microbial Community Source (Including Marine Beach Source)
| **Water Sample Type** | **Marine Beach** | **Freshwater** | **Estuarine**<br>**Sediment** | **Sediment** | **Soil** |
| --- | --- | --- | --- | --- | --- |
| **Estuary Beach** | 97% | <0.01% | 0.4% | <0.01% | <0.01% |
| **Estuary** | 94% | 2.9% | 0.2% | 0.02% | <0.01% |
| **Freshwater** | <0.01% | N/A | <0.01% | 74% | 2.6% |

**Table 1. The relative contribution of different sources to the microbial communities in** 861 **estuarine and marine water.**
| **Water Sample Type** | **Marine Beach** | **Freshwater** | **Estuarine**<br>**Sediment** | **Sediment** | **Soil** |
| --- | --- | --- | --- | --- | --- |
| **Estuary Beach** | N/A | 66% | 12% | <0.01% | <0.01% |
| **Estuary** | N/A | 74% | 7.6% | 0.02% | <0.01% |
| **Marine Beach** | N/A | 35% | 19% | <0.01% | <0.01% |
| **Freshwater** | N/A | N/A | 0% | 74% | 2.6% |

### Relationships between environmental conditions, fecal source concentrations, environmental sources and enterococci concentrations

Two PLSR models were created to determine relationships between enterococci and fecal source concentrations, environmental sources, and environmental conditions (outlined in the Methods). The first ‘freshwater’ PLSR model indicated particle-associated enterococci concentration, concentration of mammal fecal marker, TSS concentration, percent of sediment source, percent of unknown source, and salinity were important variables (VIP > 0.8) in resolving variation in enterococci concentrations (Table 1). A one-factor (single PLSR regression) model was deemed optimal (root mean PRESS = 0.735), and showed that all variables (except salinity) had positive associations with enterococci concentrations. Values for model performance (R^2^Y = 0.6, R^2^X = 0.5, and Q^2^ = 0.4) indicated that the model fit the data moderately well (R^2^X ≥ 0.5) but had poor predictive capability of enterococci concentrations (Q^2^ < 0.5; Supplementary Material 3). Out of all the important variables, particle-associated enterococci (Particle ENT) concentrations showed the strongest relationship to total enterococci concentrations (Table 2). The second PLSR model, a two-factor/two PLSR regressions model, was the best fit (root mean PRESS = 0.744) from the PLSR constructed for the estuary, estuary beach, and marine beach sites. The analysis identified particle-associated enterococci concentration, mammal fecal source concentration, percent of freshwater, unidentified and estuarine sediment sources, water temperature, and high tide height as significantly related to enterococci concentrations. Factor one showed that all variables were positively associated, except for the percent unidentified and marine sediment sources. The second factor showed mammal fecal sources, freshwater sources, and water temperatures were negatively related to enterococci concentrations, which was the opposite of their associations for factor one. The high tide height and marine sediment were positively related to enterococci concentrations for factor 2 of the PLSR (Table 2). Together both factors explained 61.8% in the variation observed in enterococci concentrations, and model performance (R^2^Y = 0.6, R^2^X = 0.5, and Q^2^ = 0.6) indicated better predictive ability with a similar fit to the data compared to the freshwater model (Supplementary Table 3). Out of all the potential variables measured (19 total) across three categories (fecal source input, environmental source contribution, and environmental conditions), particle-associated enterococci and mammal fecal marker concentrations had the most significant relationships to enterococci concentrations. The relationships between other variables and enterococci concentrations were specific to freshwater and estuary/marine beach models, indicating ecosystem specific relationships. However, the joint relationship of particle-associated and mammal fecal marker across freshwater and estuary/marine environments indicate their overarching importance in determining enterococci concentrations.

**Table 2.** Most Significant Relationships/Contributions for All Factors to Enterococci Concentrations. Shown is the output from a partial least squares regression for a freshwater and estuary/marine model. All variables shown have significant relationships for each model (VIP > 0.8), and loadings are derived from re-running models with only variables deemed significant. Model loadings are specific weights on a multivariate regression axis, positive and negative loadings refer to positive or negative relationships to enterococci concentrations. Negative loadings in the model are designated with a – before the number.

| Freshwater |  | Estuary, Estuary Beach & Marine Beach |  |  |  |
| --- | --- | --- | --- | --- | --- |
| PLSR 1 |  | PLSR 1 |  | PLSR 2 |  |
| X Variable | Loading | X Variable | Loading | X Variable | Loading |
| Particle ENT | 0.501 | Particle ENT | 0.456 | Particle ENT | 0.420 |
| qPCR Mammal | 0.352 | qPCR Mammal | 0.438 | qPCR Mammal | -0.337 |
| TSS | 0.408 | % Freshwater | 0.408 | % Freshwater | -0.418 |
| % Sediment | 0.336 | % Unknown | -0.457 | % Unknown | 0.389 |
| % Unknown | 0.476 | Water Temp (C) | 0.302 | Water Temp (C) | -0.123 |
| Salinity | -0.344 | Hightide (ft) | 0.170 | Hightide (ft) | 0.456 |
|  |  | % Estuarine Sediment | -0.294 | % Estuarine Sediment | 0.401 |
| Total Y Variance | 60.1% | Total Y Variance | 47.2% | Cumulative Y Variance | 61.8% |

## 4 Discussion

Geometric mean enterococci concentrations at the marine beach, estuary, and estuary beach sampling sites were all less than the State of Maine water quality standard of 35 CFU/100 ml and the majority of concentrations were less than the 104 CFU/100 ml single sample standard, indicating the water quality was typically considered acceptable for recreational use. Previous monitoring by the Maine Healthy Beaches Program in 2014 had shown the Wells Beach area was one of 7 beaches in Maine that had a greater than 20% exceedance rate, with suspicion that freshwater inputs are a significant source of contamination (38). Our findings confirmed that enterococci concentrations were statistically higher at both major freshwater tributaries to the estuary, especially at the Depot Brook site where levels were regularly above the 104 CFU/100 ml single sample standard. The Depot Brook site is located in a watershed with a higher fraction of developed land (0.27-0.50) and more people per km^2^ (325-2,650 people) compared to the Webhannet site watershed that has a lower developed fraction (0.13-0.25) and 150-325 people per km^2^; 40). This could help explain the difference in enterococci concentrations between freshwater sites as a more urbanized watershed can increase transport of more pollution from the watershed to the freshwater tributary. However, the summer of 2016 was especially dry in this region (41) with just one event with >1 inch of rain (1.73 in., 6/28/16) 48 h prior to the sampling time. This overall dry condition likely contributed to less fecal contamination transport (via freshwater discharge) from the watershed to the estuary and marine beach. This suggests that more typical rainfall conditions would probably have resulted in more freshwater discharge and higher enterococci concentrations than what we observed.

Enterococci were significantly associated with suspended particles of >3.0 μm diameter (R^2^ = 0.96, p < 0.05). On average, 36% (SD ± 30) of the total enterococci concentrations were associated with particles, which suggests particles as a potentially important transport mechanism. Other studies conducted in estuary and storm waters have found similar fractions of particle associated enterococci, but they noted enterococci demonstrated a preference for a larger particle size of >30 μm (42–44). The large standard deviation for particle-associated enterococci could be attributed to the complex nature of particle interactions (sedimentation rate, electrostatic, hydrophobic, and other surface-surface interactions) and hydrogeological dynamics (salinity-driven turbidity maximum) (45). The mechanisms underlying enterococci-particle interactions may also be related to ionic strength in surface waters, as *Enterococcus faecalis* is negatively charged over a broad pH range (2-8 pH units) and in the presence of different ion concentrations (46). Results for this study indicate that TSS and particle-associated enterococci had no linear relationship, indicating particle-associated enterococci were not dependent on the total amount of suspended material and thus the association is likely due to other factors influencing cell-particle interactions.

Quantitative PCR assessment of several fecal sources is a potentially useful strategy to determine the relative significance of the different sources in a single sample and over time at sites of interest. PCR detection showed a chronic presence of mammalian fecal source(s) (100% of samples) with human fecal source(s) detected in approximately half of all samples, so qPCR analysis is useful for bringing context to the significance of these findings. For example, Mayer et al. (47) showed that wastewater effluent contains about 10^8^ copies/100 ml of the AllBac mammal fecal marker, Sowah et al. (48) found that streams impacted by septic systems could contain 10^5^ – 10^7^ copies/100 ml depending on the season, and Bushon et al. (49) determined that under storm flow conditions in an urban watershed mammal marker copy numbers could exceed 10^8^ copies/100 ml. Results for this study ranged from 10^5^ to 8.6 × 10^7^ copies/100 ml, values that are within previously reported ranges and likely a concentration reflective of a predominantly non-urbanized watershed and intermediate mammal source loading. The estuary and estuary beach area showed a statistically higher concentration of the mammal marker, however, there was no responsive increase in the concentrations of the human associated fecal marker (HF183), which may indicate that humans are not the primary mammalian source for the increased fecal contamination.

The average concentration of the human marker was 1,500 copies/100 ml across all sites (geometric mean 167 copies/100 ml), with the highest concentration being 20,364 copies/100ml (Webhannet 6/22/16). Boehm et al. (50) showed that 4,200 copies/100 ml of HF183 is the cutoff for where GI illnesses exceed the EPA acceptable risk level of approximately 30/1000 for swimmers (1). On average, sites in this study did not exceed this benchmark level, however, there were 10 occasions when sites were above the 4,200/100ml threshold (7 different sites across 4 sampling dates), indicating that sporadic events or conditions can cause elevated human fecal contamination and potential public health concerns (Supplementary 4). Boehm et al. also showed that at the LOQ for most assays, 500 copies/100ml or 1000 copies/100ml, there is still a predicted GI illness of 4 or 8 cases per 1000 swimmers, suggesting positive detection at the LOQ is indicative of low level health risk (50). For this study, the LOQ was 250 copies/100ml for the HF183 assay and 67 of 117 samples (57%) tested positive at or above this limit, suggesting that over half of collected water samples indicated the presence of a low-level health risk. Although there were no statistical differences between sites for human fecal contamination, W11 did contain the highest geometric mean (493 copies/100 ml; Supplementary 4). This could be reflective of the location of the site as it’s where drainage from the Webhannet and Depot watershed meets and is also directly downstream from a boat marina with the harbor sewage pump station, which could be a possible point source of contamination. Nonetheless, even though sites on average were below published thresholds, detection of human contamination even at low concentrations is a concern.

Although human fecal sources are the greatest public health concern (6, 7, 22, 51) we did not observe any relationship between human fecal contamination and enterococci concentrations, suggesting other mammalian fecal sources are more influential in explaining the variation observed in this study. Interestingly gull fecal sources were detected in 77% or more of the samples in the estuary and marine beach area, however only 10% of the samples were positive within the fresh water (Supplementary Material 1), despite there being no decrease in the bird fecal marker concentration, suggesting the presence of different bird sources in these areas.

Anecdotally, Canada geese were observed upstream of both the Webhannet and Depot freshwater sites periodically throughout the season, which could be a significant source of bird fecal contamination in the fresh water locations (52).

One of the unique findings of this study was the relative contribution of different sources to the bacterial community in the estuarine water. The bacterial community in estuarine water primarily originated (>90%) from marine beach water, which is not surprising for a well-flushed estuary like the study site. Because the study period was minimally influenced by rainfall and associated runoff of freshwater, we expected that the influence of freshwater sources would be low. In ensuing analyses, we chose not to include marine beach water as a potential source for a variety of reasons. First, the samples were always collected during low tide before the ebb when the estuary water was draining and water was moving from the watershed towards the marine beach. Secondly, we had already shown that the OTU compositions for the marine beach and estuary samples were very similar, increasing the possibility of a type I error (false positive) for identifying marine beach as the likely source of enterococci. Lastly, fecal pollution sources most likely come from the watersheds and not from marine water, so excluding marine beach water helps to enhance the determination of watershed influences. Our second analysis (marine beach source excluded) showed that freshwater was a significant source of bacteria to the estuary (>65% assignment) compared to soil, sediment, and estuarine sediment. This implicates freshwater as a major conduit for bacterial transport, as well as the major source of enterococci to the estuary. Overall this finding highlights the importance of freshwater discharge as a controlling factor in transporting contamination from the watershed to the coast. The specific percent assignment of freshwater source could be an over-estimate, however the trend observed is a likely scenario given the rational discussed.

Analysis of environmental reservoirs of enterococci (soil, sediment, etc.) and their presence within water samples using SourceTracker revealed a variety of source contributions to freshwater, estuary and marine waters. To date there have been limited studies using SourceTracker to identify soil and sediment-associated taxa within water samples, and none of these studies have focused on a coastal watershed with the potential for freshwater, estuarine and marine sources. One study conducted in the upper Mississippi River identified up to 14% of sediment and 1.4% of soil sources of the taxa within the river water (53). This study, however showed that the sediment source was much more abundant in freshwater (74%), indicating a greater degree of mixing between the freshwater and underlying sediment communities. The amount of sediment and soil sources within water samples may be related to site specific characteristics such as relief or soil texture, which has been shown with TSS fluxes on a global scale (54). Thus, the degree to which the underlying sediment community mixes with the overlaying water is likely site specific. Interestingly, even though freshwater contained a significant amount of sediment source taxa, no sediment source was observed at the estuary and marine beach sites through the SourceTracker analysis. This difference could indicate that rapid sedimentation happens during transit to and within the estuary and at the estuarine turbidity maximum zone (55). TSS concentrations and the ratio of particle-associated to total enterococci concentrations, however, showed no differences between freshwater and estuary/marine sites. This could be related to the separate and quite different hydrodynamics within these different water systems. The percent of sediment source in the freshwater samples observed here might also be an over-estimate/over fit from SourceTracker given the limited number of potential sources used, but results consistently showed an elevated presence of sediment in all freshwater samples in this study. SourceTracker analysis also revealed that the freshwater source was significant (35% or more) in estuary and marine beach water samples, suggesting that fresh water is a significant conduit for microbial, and fecal contamination, transport from the watershed to the estuary and marine beach.

The use of predictive models for water quality has been a focus in the field in parallel with the adoption of bacterial indicator organisms as the gold standard for water quality determination. The goal of this research was to identify significant influences on enterococci concentrations by measuring a wide variety of variables. To distill this information, we used a PSLR model, which has been shown to out-perform similar multiple linear regression and principle components regression analyses (56) and has gained popularity in the water quality field (57, 58). Results from the PLSR analysis in this study showed that particle-associated enterococci and concentrations of mammal fecal sources were the driving force behind variation in enterococci concentrations, as described by both PLSR models constructed. Other factors were found to influence enterococci concentrations, however, these differed between the freshwater and estuary/marine beach models. For example, TSS concentration as well as the percent of both freshwater sediment and unknown sources positively influenced enterococci concentrations at freshwater sites. This indicates that sediment is a likely source of enterococci that influences concentrations measured in the water. Positive influences from the unidentified source taxa suggests that there is either an alternative source (not measured in this study) within the watershed that also influences enterococci concentrations or that SourceTracker could simply not resolve all the potential sources we used. This finding is not surprising given the vast number of potential sources of fecal pollution within a watershed and that fecal sources were not a part of the SourceTracker analysis. Results from the estuary and marine beach model returned a two-factor regression, with each factor essentially being the inverse of each other. Specifically, it highlighted freshwater being a major conduit for microbial transport to and through the estuary. Negative influences from the unknown source reaffirms this finding, along with positive influences from the previous high tide height. The second factor explained approximately 15% of the variation in enterococci concentration, therefore its importance must be weighed proportionately to factor one, which explained almost 50% of the variation. However, positive loadings from previous high tide height and percent of estuarine sediment indicate estuarine sediment could be a source of enterococci whose influence is dependent on tide height. The negative loadings from mammal fecal source(s) may indicate that enterococci originating from the estuarine sediment are not from mammal fecal sources.

Overall, the results from this study demonstrated that concentrations of enterococci in the coastal estuarine/marine beach study area were largely controlled by particle-associated enterococci and mammal fecal source input. The influence of these factors is likely universal across freshwater and estuarine environments, however other ecosystem factors likely play a role as well. For freshwater portions of the coastal watershed, sediment may act as a significant enterococci reservoir that is frequently re-suspended within the water column. Freshwater itself could act as a major conduit for bacterial transport to an estuary and marine beach area where other environmental factors (water temperature and high tide height) can influence enterococci concentrations as well. These findings highlight the dynamic nature of enterococci in natural aquatic ecosystems outside of the mammalian fecal tract, and that concentrations within fresh water and estuary/marine beach water are influenced by a variety of factors.

## Materials and Methods

### Site description

This study was conducted in Wells, Maine, USA (Figure 1). Eight different sites were used to monitor water quality (n = 2 freshwater, n = 2 estuary, n = 3 estuary beaches, n = 1 marine beach) as well as twelve soil, twelve fresh-water sediment and four estuarine sediment sampling sites. Data for air temperature and rainfall amount for the 48 h prior to sampling were obtained from Weather Underground (https://www.wunderground.com/cgibin/findweather/getForecast?query=Wells,%20ME) and characteristics of tides during sampling were obtained from US Harbors (www.meusharbors.com).

### Water sampling

Surface water samples were collected weekly from June to September 2017 (n = 117). Sampling started two hours before low tide to maximize the potential impacts of freshwater pollution sources, and samples from all estuary and marine beach sites were collected before the slack tide. Water samples were collected in autoclaved 1L Nalgene™ Wide-Mouth Lab Quality PPCO bottles (Thermo Fisher Scientific, Waltham, MA, USA), and environmental parameters were measured with a YSI Pro2030^®^ dissolved oxygen, conductivity, and salinity Instrument (YSI Incorporated, Yellow Springs, Ohio, USA). A field replicate was collected at a different site for each sampling event.

### Soil, sediment, and marine sediment collection

Environmental sources were collected twice throughout the sampling season to build source libraries that were “finger-printed” with 16S sequencing and SourceTracker analysis. Six soil and sediment samples were collected upstream of both freshwater sites (Webhannet and Depot; Figure 1). Soil samples were collected at the crest of the stream embankment, where a 10 × 10 cm a plastic square template was placed down and all soil (O-horizon) within the template at a 2 cm depth was collected. Samples were sieved (USA Standard No. 5) to remove any loose-leaf litter and roots to only sample smaller soil particles and their microbes. Underlying stream sediments were collected using a Van Veen sediment sampler from depositional sites chosen based on the presence of fine grain sediments. One grab sample was collected for each site and then the top 2 cm of sediment was subsampled for analysis. Sediments were sieved (USA Standard No. 45) to remove coarse grain and gravel size particles. Estuarine sediments were collected during low tide when intertidal sediments were exposed using the Van Veen sampler, and the top 2 cm were again collected for analysis.

### Enterococci and total suspended solids quantification

Total and particle-associated enterococci were enumerated using the EPA Method 1600 membrane filtration protocol (59) and particle-associated enterococci were determined via filtration through a 0.47 mm diameter 3.0 μm pore size polycarbonate filter (Millipore™, Darmstadt, Germany) as first reported by Crump et al. (60). The filters were rolled onto plates containing mEI agar and incubated at 41°C ± 0. 5°C; representative colonies were counted in 24 ± 2 hours. Total suspended solids (TSS) were measured using EPA method 160-2, where 500 ml of the water sample was used to determine TSS concentrations (61).

### DNA extractions

DNA extraction from all matrices was performed with the PowerSoils^®^ DNA Extraction Kits (MO BIO Laboratories, Carlsbad, CA, USA), with modifications to the manufacture’s protocol needed to optimize the extraction from water sample filters. For water samples, 500 ml collected water sample was filtered through 0.47 mm diameter 0.45 μm pore size polycarbonate filter (Millipore™, Darmstadt, Germany), which was stored in a sterile 2 ml cryotube at -80°C for at least 24 h. Prior to DNA extraction, frozen filters were crushed into small pieces with an ethanol sterilized razor blade, a practice commonly used to maximize DNA recovery (62–64). To minimize additional DNA loss during the extraction process solutions C2 and C3 (from manufacturer’s protocol) were halved in volume and combined into a single step. DNA extraction from soil, freshwater sediment, and marine sediment were conducted per the manufacture’s protocol.

### Microbial source tracking (MST) PCR and qPCR assays

MST PCR assays that target Mammals (Bac32; 65), Humans (HF183; 9), Gulls (Gull2; 66), Dogs (DF475; 10) and Ruminants (CF128; 9) were used to determine the presence of fecal sources in water samples. Positive control plasmids were created for each PCR assay from fresh fecal samples that came from each target organism (Human, Gull, Dog, and Cow). The TOPO TA Cloning Kit was used (Invitrogen, Carlsbad, CA, USA), with a blue/white screen of *E. coli* transformants on kanamycin (50 μg/mL) selective TSA plates. Positive *E. coli* colonies were screened with their respective PCR assay, and PCR positive colonies were then grown in TSB and extracted with the PureLink^®^ Quick Plasmid Miniprep Kit (Invitrogen, Carlsbad, CA, USA). PCR assays were run on a T100™ Thermal Cycler (BioRad, Hercules, CA, USA) with the GoTaq^®^ Green MasterMix (Promega, Madison, WI, USA). Cycling conditions and amplification protocols for each assay targeted the different source specific markers and followed protocols delineated by different studies: Bac32 (67) and HF183 (67), CF128 (68), DF475 (69), and Gull2 (66). Quantitative PCR assays were also run to determine fecal source strength for Mammals (AllBac; 70), Humans (HF183; 71), and Birds (GFD; 72). All qPCR assays were run on a Mx3000P cycler (Agilent Technologies, Santa Clara, CA, USA), TaqMan assays used the PerfecCTa^®^ FastMix^®^ II (QuantaBio, Beverly, MA, USA) master mix and the SYBR green assay used the FastSYBR™ Green Master Mix (Applied Biosystems, Foster City, CA, USA). A standard curve ranging from 10^6^–10^2^ copies (Mammal assay) or 10^5^–10^1^ copies (Human & Bird assay) was also run for each experimental run with the limit of quantification (LOQ) being 100 copies (Mammal) or 10 copies (Human & Bird) per PCR. The Ct values, amplification efficiency, slope, and R^2^ values for each standard curve were compared to previously run standard curves, to ensure satisfactory performance before being used to calculate copy numbers for that run. Each environmental sample was diluted 1:10 and run in triplicate and the reaction volume (25 μl) contained a final concentration of 0.2 mg/ml BSA. Amplification/cycling conditions were preformed per published protocols for AllBac (73), HF183 (73), and GFD (16). TaqMan assays were run with an internal amplification control (74) with a down-shift of 1 cycle considered inhibition. Samples spiked with a plasmid containing 10^4^ copies of GFD amplicon were used as inhibition controls for the SYBR assay, with a recovery of less than 10^4^ copies (100%) considered inhibition. For a list of primers, probes, and standard curve performance, see Supplementary Material 1.

### 16S library preparation

The V4 region of the 16S rRNA gene, using the 515F-806R primer-barcode pairs, was used for amplicon sequencing (75). The Earth Microbiome Project protocol was used for amplification and pooling of samples, with minor modifications (76). The Qubit^®^ dsDNA HS assay was used to quantify sample concentrations, and 500 ng of DNA was pooled per sample. The pool was then run on a 1.2 % low-melt agarose gel to separate primer-dimers from acceptable product, and bands between 300-350 bps were cut and extracted as described above. The final DNA sample was then run on the Agilent Technologies 2200 TapeStation system (Santa Clara, CA, USA) to determine final size, quality, and purity of sample. Each library was sent to the Hubbard Center for Genome Studies at the University of New Hampshire to be sequenced (2 × 250 bp) on the Illumina HiSeq 2500 (San Diego, CA, USA).

### Quality filtering and Operational Taxonomic Unit (OTU) picking

QIIME 1.9.1 was used to perform all major quality filtering, and OTU picking (77). Forward and reversed reads were quality trimmed (μ P25) and removed of Illumina adapters via Trimmomatic (78). Any reads that were less than 200 bps were discarded, and reads were merged with the QIIME joined_paired_ends.py, using a minimum overlap of 10 bps and a maximum percent difference of 10%. Paired-end data were analyzed using the QIIME open-reference OTU picking strategy with UCLUST for *de novo* picking and the Greengenes 13_8 database (79) for taxonomic assignment. Alternative OTU picking strategies were also tested to determine best workflow, for performance of difference strategies refer to Supplementary Material 2. Data for all sequenced samples are publicly available through NCBI BioProject (http://www.ncbi.nlm.nih.gov/bioproject/431501).

### SourceTracker analysis

Samples from 4 source types (fresh water, soil, sediment, and marine sediment) and 4 sink types (fresh water, estuary water, estuary beach water, and marine beach water) were analyzed by the open-source software SourceTracker v1.0 (37). Default parameters were used (rarefaction depth 1000, burn-in 100, restart 10, alpha (0.001) and beta (0.01) dirichlet hyperparameters) in accordance with previously published literature (53, 80). A ‘leave one out’ cross validation was performed to assess the general performance of the model and source samples were iteratively assigned as sinks to assess how well a known sink would be assigned (i.e. source = soil and sink = soil). The percent assignments from SourceTracker are the result of the Gibbs Sampler assigning OTUs from an unknown sample to sources in a random and iterative fashion, and then calculating likelihood of that OTU originating from said source. The final output can be interpreted as the percent (or likelihood) of OTUs present in an unknown sample originating from the sources used in the analysis

### Partial least squares regression model

A partial least squares regression (PLSR) model was used to determine the most important and significant variables affecting enterococci concentrations (81). Two models were created, one for the estuary, estuary beach, and marine beach sites, and one for the freshwater sites. Particle-associated enterococci, environment variables (water temperature, air temperature, dissolved oxygen, salinity, height of previous high tide, rainfall in previous 48 h), fecal source strength (mammal, human, and bird), and percent of environmental source (fresh water, soil, sediment, and marine sediment) were used as explanatory variables for the non-freshwater model. The same parameters, except height of previous high tide and percent of freshwater source, were used for the freshwater model. All data except the percent assignments from SourceTracker were log (x+1) transformed before performing the analysis. A KFold cross validation (K=7) with the NIPALS method was used to determine optimal factors and variable importance (VIP > 0.8) for each model. Models were then re-run with only explanatory variables that were determined to be significant. To see model validation and diagnostic plots, refer to Supplementary Material 3.

### Routine statistical analysis and data visualizations

All routine statistical analyses were performed in R v3.4.0, Python 3.6.1, or JMP Pro13, while multivariate analyses were performed with PC-ORD v6. Graphing was performed in IPython notebook with matplotlib, seaborn, pandas, and numpy packages. All pairwise comparisons were done using the Kruskal-Wallis nonparametric method, with Dunn’s nonparametric multiple comparisons run *post hoc* using a Bonferroni correction.

## ACKNOWLEDGEMENTS

We would like to thank Meagan Sims and Keri Kaczor at the Maine Healthy Beaches program and Sean Smith PhD at the University of Maine for their guidance and help with general knowledge of the Wells, ME area and planning of field sampling. Field work, sample processing, and molecular work were assisted by Christine Bunyon, Alexandra Bunda, Jackie Lemaire, Audrey Beresnson, and Joseph Sevigny assisted in optimizing bioinformatic workflows. This work was funded by the National Science Foundation New Hampshire EPSCoR IIA-1330641 grant.

